# R-Loop Functions in *Brca1*-Associated Mammary Tumorigenesis

**DOI:** 10.1101/2024.02.14.580374

**Authors:** Huai-Chin Chiang, Leilei Qi, Payal Mitra, Yanfen Hu, Rong Li

## Abstract

Excessive R-loops, a DNA-RNA hybrid structure, are associated with genome instability and *BRCA1* mutation-related breast cancer. Yet the causality of R-loops in tumorigenesis remains unclear. Here we show that R-loop removal by *Rnaseh1* overexpression (Rh1-OE) in *Brca1*-knockout (BKO) mouse mammary epithelium exacerbates DNA replication stress without affecting homology-directed DNA repair. R-loop removal also diminishes luminal progenitors, the cell of origin for estrogen receptor α (ERα)-negative BKO tumors. However, R-loop reduction does not dampen spontaneous BKO tumor incidence. Rather, it gives rise to a significant percentage of ERα-expressing BKO tumors. Thus, R-loops reshape mammary tumor subtype rather than promoting tumorigenesis.

## Introduction

R-loops, a three-stranded DNA-RNA hybrid structure, consists of an RNA strand annealed to one strand of a double-stranded DNA molecule and the complementary single-stranded DNA. R-loops are formed during transcription, DNA replication, and DNA repair in both prokaryotic and eukaryotic genomes (Brickner et al. 2022; Petermann et al. 2022). A wealth of evidence implicates R-loops in diverse physiological processes including transcriptional activation and repression (Boque-Sastre et al. 2015; Grunseich et al. 2018), DNA replication (Aguilera and Garcia-Muse 2012), immunoglobulin class switch recombination (Yu et al. 2003), CRISPR-Cas9-mediated DNA editing (Jinek et al. 2012), and homology-directed repair (HDR) of double strand DNA breaks (DSBs) (Ohle et al. 2016; Ouyang et al. 2021). In addition, R-loops have also been implicated in relieving DNA topological stress (Chedin and Benham 2020), which otherwise could lead to DNA breaks (Bacolla and Wells 2004; Zhao et al. 2010). In contrast to the regulatory functions of R-loops in various DNA transactions, unscheduled R-loop formation is a significant contributor to DNA replication stress, replication-independent DNA damage, and ultimately genome instability (Aguilera and Garcia-Muse 2012; Skourti-Stathaki and Proudfoot 2014). Accordingly, a growing number of proteins have been identified for their roles in prevention and/or resolution of unscheduled R-loops (Brickner et al. 2022; Petermann et al. 2022). Indeed, inactivation of these proteins is associated with aberrant R-loop accumulation in various human diseases including cancers (also see below). Because R-loops are a well-documented source of genome instability, they are commonly presumed to serve as a driving force in tumorigenesis. However, to date there is no direct evidence for a causal role of R-loops in cancer development.

Women with certain heterozygous germline *BRCA1* mutations (*BRCA1^mut/+^*) have up to 80% lifetime risk of developing breast cancer (Kuchenbaecker et al. 2017). *BRCA1*-associated breast tumors tend to be basal-like and lack the expression of estrogen receptor α (ERα), progesterone receptor, and HER2 (so called “triple-negative”)(Visvader and Stingl 2014). Multiple lines of evidence strongly indicate that luminal progenitor cells of *BRCA1^mut/+^* breast tissue serve as the cell of origin that gives rise to *BRCA1*-associated breast tumors (Lim et al. 2009; Molyneux et al. 2010; Proia et al. 2011). These luminal progenitor cells from precancerous *BRCA1^mut/+^* breast tissue are defective in differentiation into mature luminal cells (Lim et al. 2009). Furthermore, the expression of luminal differentiation genes is significantly reduced in *BRCA1^mut/+^*breast epithelium versus non-mutation carriers (Proia et al. 2011). More recent studies suggest that these differentially blocked *BRCA1^mut/+^* luminal progenitor cells undergo oncogenesis upon further stimulation by hormonal and DNA damage-induced signals (Nolan et al. 2017).

At the molecular level, the BRCA1 protein is best known for its role in HDR of DSBs and suppression of DNA replication stress (Chen et al. 2018; Venkitaraman 2019). Cell line-based studies implicate BRCA1 in R-loop elimination (Bhatia et al. 2014; Hatchi et al. 2015). BRCA1 was also shown to bind directly to DNA-RNA hybrids (D’Alessandro et al. 2018), raising the possibility that BRCA1 may play a role in sensing R-loops. Using clinical samples and mouse models, our published work showed significant accumulation of R-loops in luminal epithelial cells harboring *BRCA1* mutations (Zhang et al. 2017). Furthermore, these *BRCA1* mutation-associated R-loops tend to be localized at transcriptional enhancers and promoters (Zhang et al. 2017). Collectively, these published studies clearly indicate a functional link between *BRCA1* mutations and aberrant R-loop accumulation. The prevailing paradigm is that R-loops and their associated genome instability directly lead to *BRCA1* mutation-associated tumor incidence.

To test the hypothesis that aberrant R-loop accumulation observed in BRCA1-deficient luminal epithelial cells directly contribute to *BRCA1*-associated tumorigenesis, we established a novel transgenic mouse model whereby the mouse *Rnaseh1* gene, encoding the nuclear form of RNase H1 for R-loop dissolution (Cerritelli and Crouch 2009), is overexpressed in mammary epithelium with or without *Brca1* deletion. Contrary to the aforementioned hypothesis, mammary epithelium-specific R-loop removal did not affect the incidence of spontaneous mammary tumors from *Brca1* mutant mice. However, R-loop attenuation resets the equilibrium between luminal progenitor and mature luminal cells, resulting in a significant percentage of ERα^+^ *Brca1*-associated mammary tumors. Thus, our findings provide *in vivo* evidence for a previously unappreciated role of R-loops in *Brca1*-associated mammary tumor development.

## Results and Discussion

### RNase H1 overexpression attenuates *Brca1*-associated R-loop accumulation in mouse mammary epithelium

We previously reported substantial R-loop accumulation in mice with deletion of *Brca1* in mammary epithelium (*MMTV-Cre, Brca1^f/f^* or BKO) (Zhang et al. 2017). This mouse model, commonly used for studying *Brca1*-associated mammary tumor development (Xu et al. 1999), was chosen to interrogate a causal relationship between R-loops and tumorigenesis. To attenuate R-loop accumulation in *Brca1*-deficient mammary epithelium, we first inserted at the Rosa26 locus of the mouse genome a conditional transgenic (Tg) expression cassette for mouse *Rnaseh1*, the mRNA of which is produced only when the loxP-flanked transcription stop sequence between the promoter and the transgene is removed through the action of the Cre recombinase (fig. S1A). Through serial breeding with the *MMTV-Cre* (Wagner et al. 1997) and *Brca1^f/f^* strains (Xu et al. 1999), we generated mammary epithelium-specific *Rnaseh1* transgenic mice (*MMTV-Cre,Rnaseh1^Tg/+^* or Rh1-OE) and corresponding compound mice with both *Rnaseh1* transgene and *Brca1* deletion (*MMTV-Cre,Brca1^f/f^,Rnaseh1^Tg/+^*or BKO-Rh1-OE). Genotyping confirmed the allelic status of the floxed *Brca1*, floxed *Rnaseh1*, and Cre-encoding gene (fig. S1B).

Immunohistochemistry (IHC) confirmed that RNase H1 was overexpressed in mammary epithelium of both Rh1-OE and BKO-Rh1-OE mice (Fig. 1A). Due to the lack of a suitable mouse BRCA1-specific antibody (Yang et al. 2021), we analyzed *Brca1* mRNA levels in sorted stromal cells (EpCAM^−^CD49f^−^, Vimentin^high^), basal epithelial cells (EpCAM^med^CD49f^high^, K14^high^), and luminal epithelial cells (EpCAM^high^CD49f^med^, K18^high^) from various mouse strains (fig. S2A-B). As expected, both BKO and BKO-Rh1-OE mice exhibited substantially reduced *Brca1* mRNA levels in both basal and luminal epithelial compartments, but not in the stromal compartment (Fig. 1B). Collectively, these data ascertain cell type-specific *Brca1* gene deletion and RNase H1 overexpression.

**Figure. 1.**
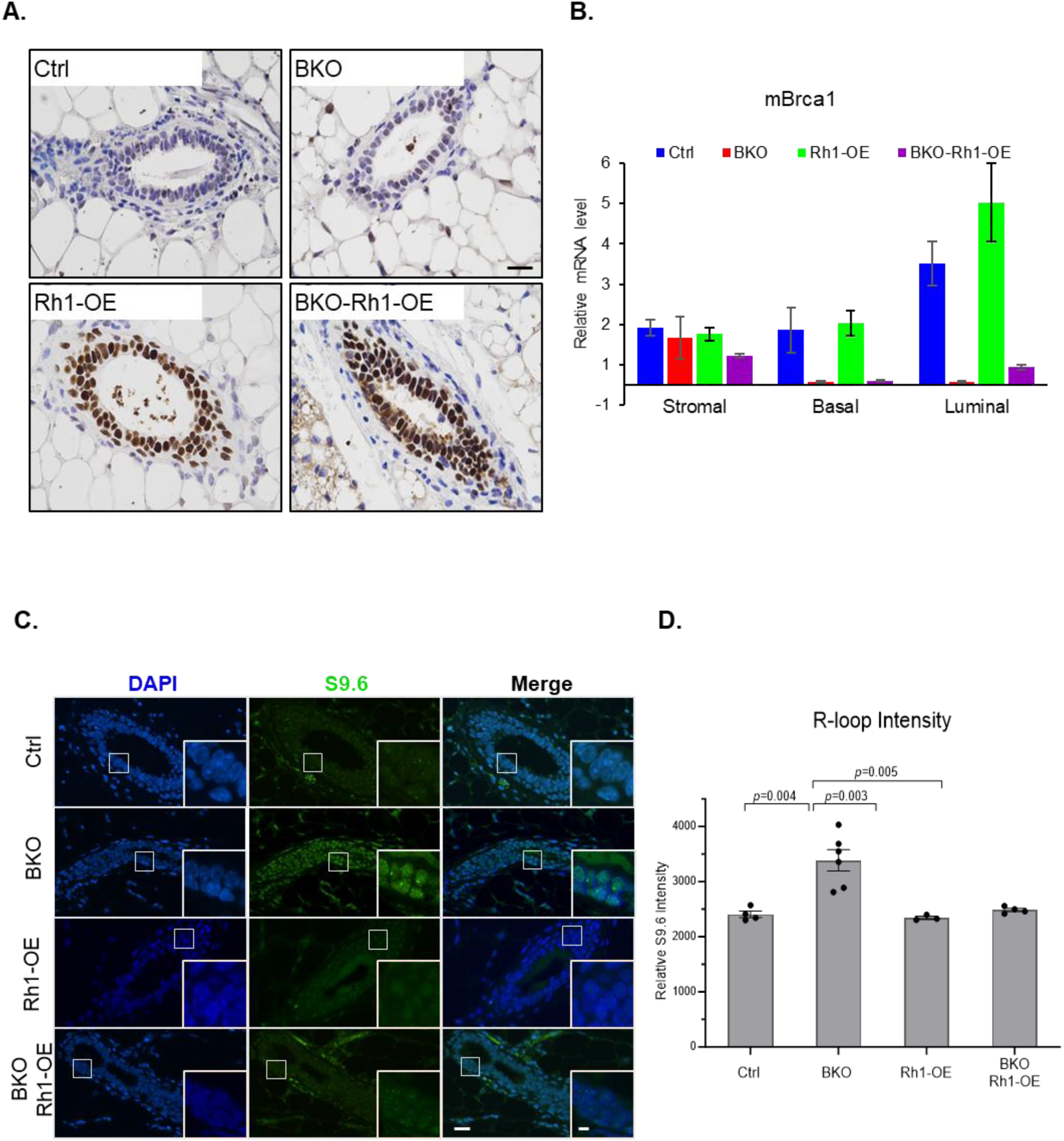
RNase H1 overexpression in the mouse mammary gland can reduce R-loop signal *in vivo*. (**A**) RNase H1 immunohistochemistry analysis in the mammary glands of 8-week virgin mice. Representative results from at least 4 sets of animals. Scale bar = 20µM. (**B**) mRNA analysis of *Brca1* using sorted stromal, basal and luminal cells. The numbers of animals used are: Ctrl = 4, BKO = 3, Rh1-OE = 3, BKO-Rh1-OE =3. Error bars represent s.e.m. (**C**) Low and high (inlet) magnification IF images of R-loop staining in the mammary ducts of 8-week-old virgin mice. Scale bars, 50 μm and 10 μm (inlet). (**D**) Quantitation of the relative R-loop intensity in 8-week-old animals. The number of animals used in each group is: Ctrl=4, BKO=6, Rh1-OE=3, and BKO-Rh1-OE=4. Statistical analysis was performed using two-tailed *t*-test. *P* value are indicated. Error bars represent s.e.m.

Next, we assessed the effect of RNase H1 overexpression on R-loop intensity in mouse mammary glands. Using the S9.6 antibody that preferentially recognizes DNA-RNA hybrids (Boguslawski et al. 1986), we detected by immunofluorescence (IF) staining prominent R-loop signals in BKO, but not control, mammary epithelium (Fig. 1C-D). This is consistent with our previously reported observation (Zhang et al. 2017). Recent studies indicate that the same antibody can bind to non-R-loop RNA structures, which could complicate IF-related data analysis and interpretation (Smolka et al. 2021). To validate the R-loop-specific IF signal in BKO mammary tissues, we pretreated them simultaneously with RNase T1 and RNase III, which specifically degrade single-stranded RNA (ssRNA) and double-stranded RNA (dsRNA), respectively (Smolka et al. 2021). The pretreatment largely eliminated cytoplasmic IF staining but retained the prominent nuclear signals (fig. S3A). In contrast, pretreatment with RNase H1 abolished the nuclear staining (fig. S3A). Together, these results confirm specificity of the S9.6 antibody in detection of the R-loop signals in *Brca1*-deficient mouse mammary epithelium. Notably, *in vivo* RNase H1 overexpression in *Brca1*-deficient mouse mammary epithelium (BKO-Rh1-OE) significantly diminished the R-loop levels compared to those in BKO (Fig. 1C-D), thus validating the expected impact of RNase H1 overexpression on R-loops *in vivo*.

### RNase H1 overexpression does not affect mammary gland development or function

Mammary epithelium-specific *Rnaseh1* transgenic mice (Rh1-OE) were born with no overt developmental defects (data not shown). The whole-mount staining of the mammary glands in virgin Rh1-OE mice and postpartum Rh1-OE mice showed normal epithelial ductal and alveolar structures, respectively (Fig. 2A-B and fig. S3B). Normal alveologenesis and lactogenesis of postpartum Rh1-OE mice were further confirmed by hematoxylin & eosin (H&E) staining (Fig. 2C) and IHC for anti-milk proteins (Fig. 2D). Moreover, pups of Rh1-OE dams were properly nursed (data not shown), again indicative of normal lactating function of the transgenic mice. Therefore, RNase H1 overexpression does not affect normal mammary gland development or function.

**Figure. 2.**
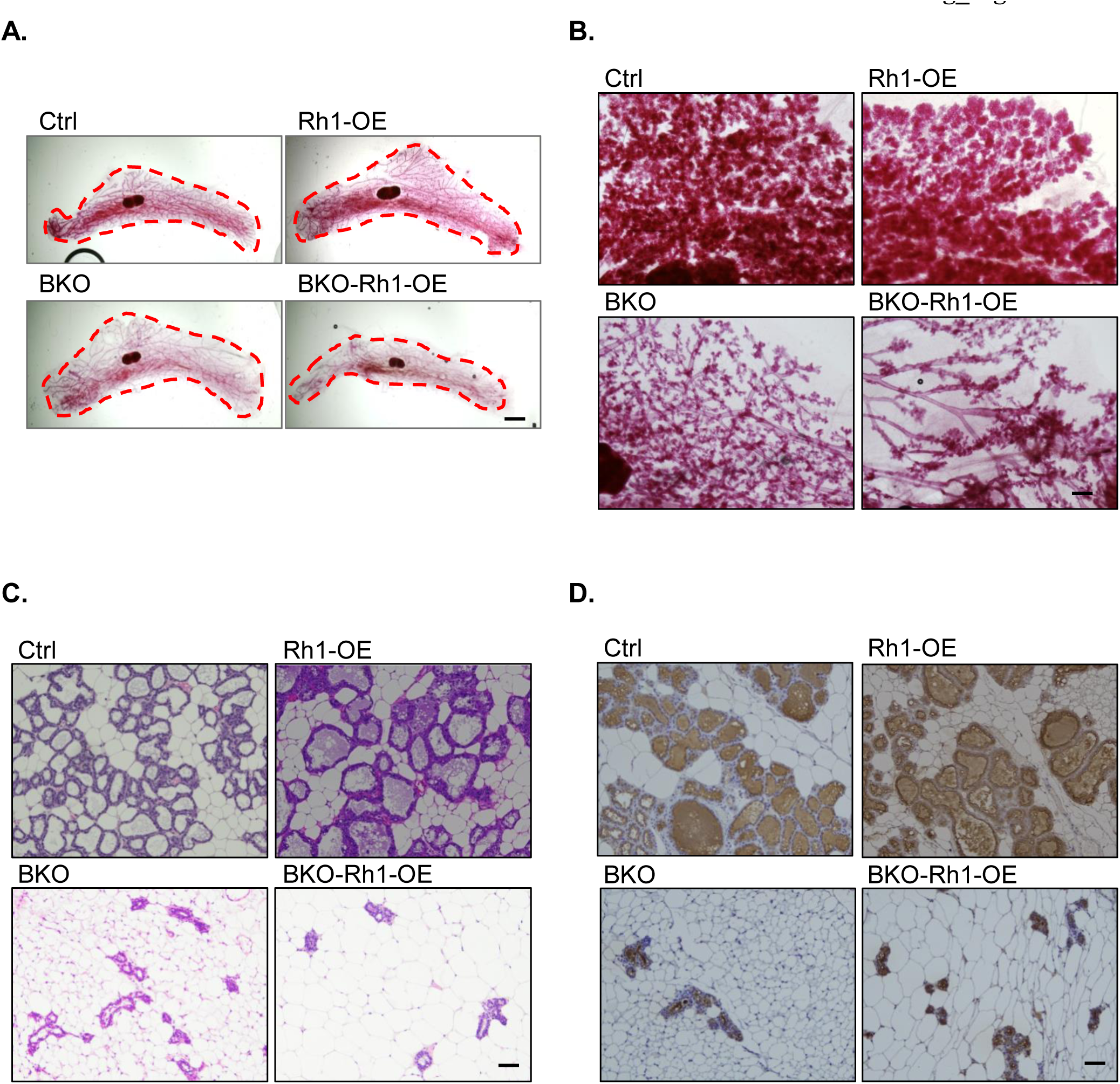
RNase H1 overexpression cannot rescue the *Brca1*-associated deficiency in alveologenesis and lactogenesis. (**A**) Whole mounts of the mammary glands from 8-week virgin mice. The red dash line highlights the boundary of the ductal area. Scale bars = 2 mm. (**B**) Whole mounts of the mammary glands from 16 to 20-week mice 1-day postpartum. Scale bar= 0.5 mm (**C**) H&E staining of the lobular-alveolar structure in mammary glands of 16–20-week mice 1-day postpartum. Scale bar= 50 μm. (**d**) IHC for total milk proteins in mammary glands of 16–20-week mice 1-day postpartum. Scale bar= 50 μm. Images in this figure are representatives of at least 4 animals in each genotype.

Consistent with published findings (Xu et al. 1999; Nair et al. 2016), the mammary glands of virgin *Brca1*-deficient mice (BKO) displayed normal mammary ductal structure (Fig. 2A and fig. S3B) but those of postpartum BKO were largely devoid of alveolar structure and milk production (Fig. 2B-D). BKO-Rh1-OE mice exhibited a similar degree of alveologenic and lactogenic deficiency (Fig. 2B-D), suggesting that RNase H1 overexpression does not rescue *Brca1*-associated defects in mammary functions.

### RNase H1 overexpression exacerbates DNA replication stress *in vivo*

To examine the impact of BRCA1 and R-loops, alone and together, on DNA replication, we pulse-labeled mice with bromodeoxyuridine (BrdU) to track cells undergoing DNA replication. Mammary tissues were immunostained for BrdU, the DSB marker γH_2_AX, and/or the HDR marker RAD51 (Fig. 3A-C, and fig. S4). As a surrogate marker for DNA replication stress, we quantified the percentage of γH_2_AX^+^ cells among BrdU^+^ mammary epithelial cells (γH_2_AX^+^/BrdU^+^) in non-irradiated animals.

**Figure. 3.**
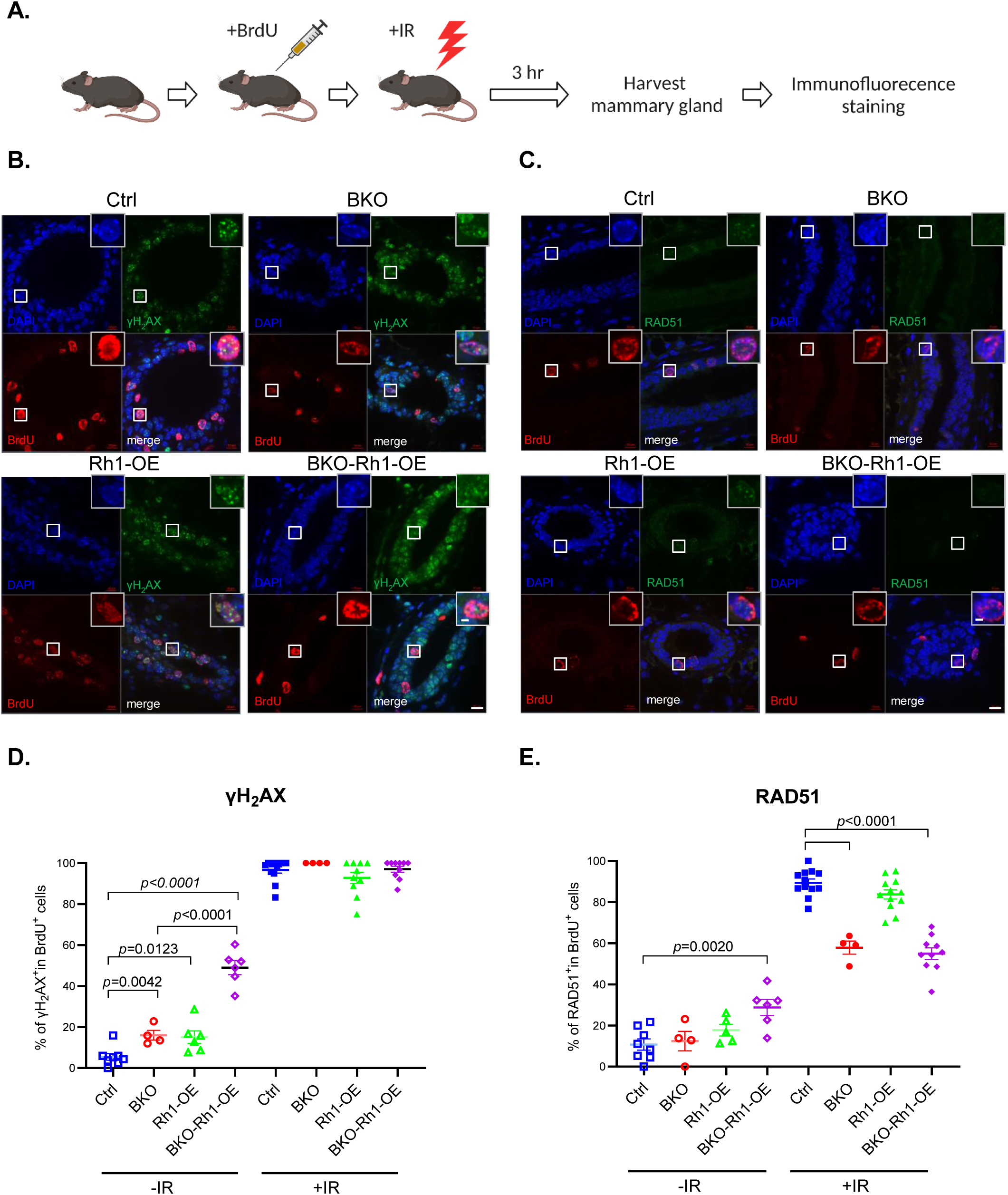
RNase H1 overexpression contributes to elevated replication stress without affecting IR-induced HDR. (**A**) Schematic illustration of the experiment setup for BrdU/γH_2_AX/RAD51 detection *in vivo*. (**B**) Low and high magnification (inlet) IF images of γH_2_AX and BrdU staining in the mammary ducts of 8-week-old virgin mice. Scale bars, 10 μm and 2 μm (inlet). (**C**) Low and high magnification (inlet) IF images of RAD51 and BrdU staining in the mammary ducts of 8-week-old virgin mice. Scale bars, 10 μm and 2 μm (inlet). (**D**) Quantification of γH_2_AX^+^/BrdU^+^ epithelial cells in the mammary gland. Each dot represents a mammary gland. Statistical analysis was performed using two-tailed *t*-test. *P* value are indicated. Error bars represent s.e.m. (**E**) Quantification of RAD51^+^/BrdU^+^ epithelial cells in the mammary glands. Each dot represents a mammary gland. Statistical analysis was performed using two-tailed *t*-test. *P* value are indicated. Error bars represent s.e.m.

As a positive control, we observed a substantial increase in DNA replication stress in non-irradiated BKO mammary glands versus their wildtype control (Ctrl) counterparts (Fig. 3D, compare columns 1 and 2). This is consistent with the known function of BRCA1 in the reduction of DNA replication stress. Given that R-loops are recognized as a source of replication stress and genome instability (Brickner et al. 2022; Petermann et al. 2022), we had anticipated that R-loop removal by *in vivo* RNase H1 overexpression would alleviate DNA replication stress. Contrary to our prediction, Rh1-OE mammary glands also displayed elevated γH_2_AX^+^/BrdU^+^ cells (Fig. 3D, column 3), which seems incongruent with the notion that R-loops contribute to DNA replication stress. Even more remarkably, compared to non-irradiated BKO and Rh1-OE mice, mammary epithelium of non-irradiated BKO-Rh1-OE mice experienced a drastic increase in the number of γH_2_AX^+^/BrdU^+^ cells (Fig. 3D, compares columns 2 and 3 with 4). This was accompanied by more RAD51^+^/BrdU^+^ cells in non-irradiated BKO-Rh1-OE mice compared to the Ctrl, BKO and Rh1-OE groups (Fig. 3E, compare columns 1-3 with 4), suggesting that the elevated replication stress is unlikely due to impaired RAD51 recruitment to stalled replication forks. We infer from these results that R-loop elimination and *Brca1* deletion act concertedly to accentuate DNA replication stress in proliferating mammary epithelial cells.

BRCA1 is known to play an important role in protection of nascent DNA strands at stalled DNA replication forks (Chen et al. 2018; Venkitaraman 2019). Specifically, a concerted action of BRCA1, BRCA2 and RAD51 protects a stalled replication fork from nuclease degradation and subsequent fork collapse (Schlacher et al. 2012; Chaudhuri et al. 2016; Kolinjivadi et al. 2017; Willis et al. 2017). We envision several scenarios whereby RNase H1 overexpression elevates replication stress in proliferating mammary epithelial cells. First, overexpressed RNase H1 could remove the small RNA-DNA hybrids formed during Okazaki fragment synthesis, disrupting normal DNA replication. Second, excessive R-loop removal may lead to a higher level of under-twisted superhelical stress throughout the genome (Chedin and Benham 2020). This, in turn, may favor the formation of non-B DNA structures such as cruciform, slipped structures, triplexes, and G-quadruplexes, which can impede DNA replication and create DNA sites susceptible to breaks (Bacolla and Wells 2004; Zhao et al. 2010). Lastly, our *in vivo* result could be explained by a potential positive role of R-loops in resolution of DNA replication stress, although such a scenario would not be compatible with the published *in vitro* data. More studies are warranted to further interrogate the R-loop function in DNA replication stress.

### RNase H1 overexpression does not significantly affect ionizing radiation (IR)-induced homology-directed repair (HDR) *in vivo*

To examine the combined impact of BRCA1 and R-loops on IR-triggered HDR in mammary epithelium, we first pulse-labeled mice with BrdU and subsequently subjected them to IR. Mammary tissues were harvested three hours following IR and immunostained for BrdU, γH_2_AX, and/or RAD51 (Fig. 3A-C, and fig. S4). We assessed the efficiency of IR-induced HDR by enumerating the percentage of RAD51^+^ cells in BrdU^+^ mammary epithelial cells (RAD51^+^/BrdU^+^).

As expected, IR-induced γH_2_AX foci were present in almost all BrdU^+^ mammary epithelial cells in the four mouse cohorts (Fig. 3D, columns 5-8). Consistent with the established role of BRCA1 in supporting HDR, irradiated BKO mammary glands exhibited a substantially lower percentage of RAD51^+^/BrdU^+^ cells versus Ctrl (Fig. 3E, compare columns 5 and 6). R-loop attenuation by RNase H1 overexpression did not significantly affect the RAD51^+^/BrdU^+^ percentage in irradiated mammary glands, either with (Rh1-OE) or without (BKO-Rh1-OE) the functional *Brca1* gene (compare column 5 and 7, 6 and 8 in Fig. 3E), although there is a trend of reduction in Rh1-OE versus parental Ctrl. Therefore, our *in vivo* data do not support an indispensable role of R-loops in HDR in mammary epithelial cells.

*In vitro* work suggests that BRCA1 is involved in multiple distinct steps of HDR: it facilitates 3’ end resection and RAD51 filament formation, an early and late step of HDR, respectively (Sy et al. 2009; Gao et al. 2014; Chen et al. 2018; D’Alessandro et al. 2018; Venkitaraman 2019). At the face value, our *in vivo* result is not consistent with such an HDR-promoting activity of R-loops. However, global R-loop removal by overexpressed RNase H1 may cancel out the HDR-promoting and -impeding activities of R-loops as previously observed in various model systems (Brickner et al. 2022; Petermann et al. 2022). In addition, we cannot exclude the possibility that HDR-associated R-loops at DSBs in mammary epithelial cells may be shielded from the action of overexpressed RNase H1 *in vivo*.

### RNase H1 overexpression reshapes *Brca1*-associated tumor subtype without affecting overall tumor incidence

To directly determine the role of R-loops in tumorigenesis, we monitored spontaneous mammary tumor development in Ctrl, BKO, Rh1-OE, and BKO-Rh1-OE female mice up to 75 weeks of age. To ensure continuing activation of the hormone-responsive promoter that drives the expression of the Cre transgene, all mice were mated throughout the entire duration of the tumor study. Consistent with the published findings (Xu et al. 1999; Zhang et al. 2017), BKO mice had an increased incidence of spontaneous mammary tumors, resulting in approximately 50% tumor-related mortality (Fig. 4A, red). In contrast, no mammary tumors were observed in mice with RNase H1 overexpression alone (Fig. 4A, green). Tumor incidence of BKO-Rh1-OE mice was indistinguishable from that in BKO (Fig. 4A, compare red and purple). Thus, despite global R-loop removal in *Brca1*-deficient mammary epithelium, the overall rate of mammary tumor development remained the same. This is not concordant with the marked increase in replication stress in BKO-Rh1-OE mice versus BKO.

**Figure 4.**
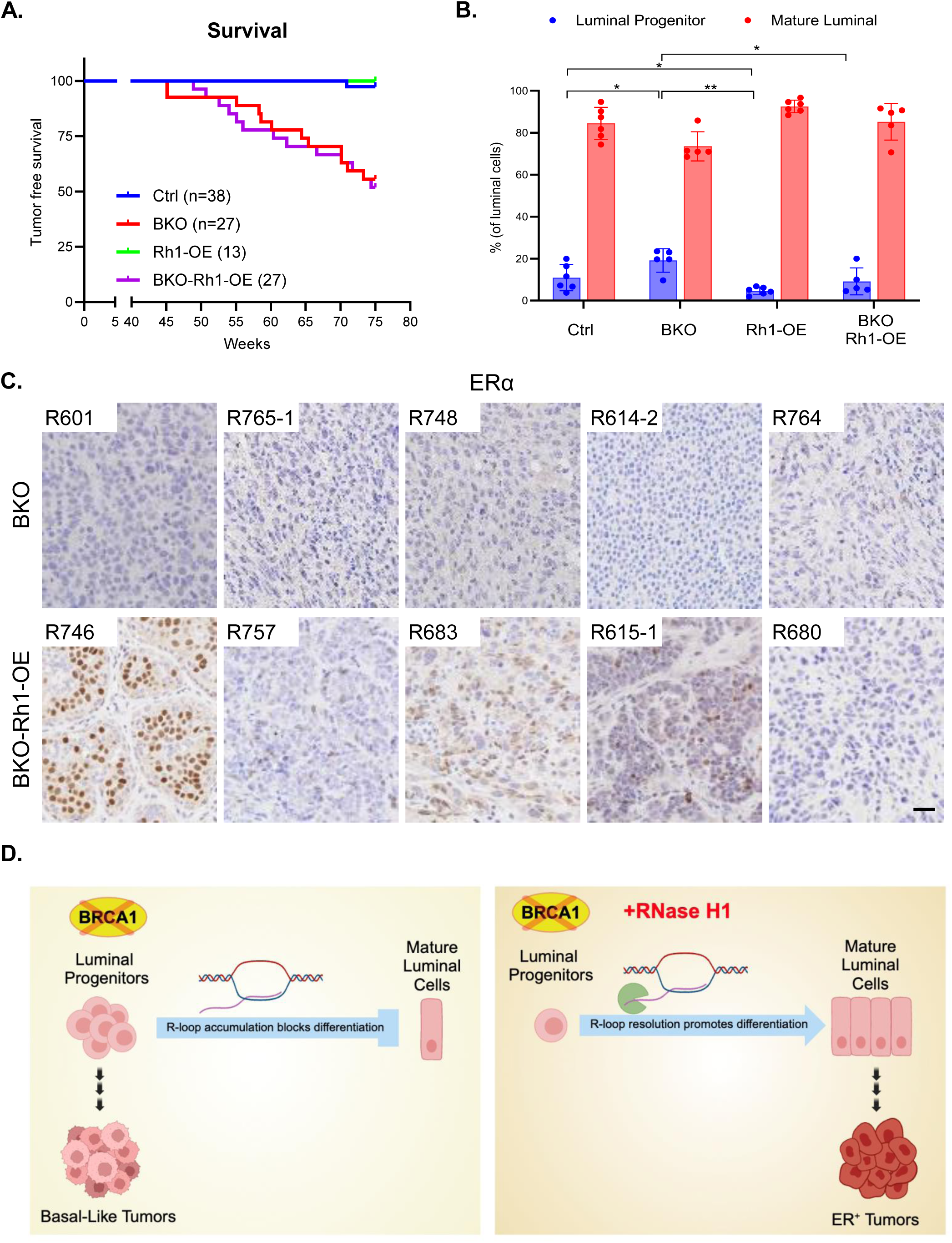
RNase H1 overexpression does not affect *Brca1*-associated tumor incidence but changes tumor subtype. (**A**) Kaplan-Meier curve for mammary tumor incidence. (**B**) Flow cytometry analysis of luminal progenitor and mature luminal cell percentages in 8-10 weeks mice. The numbers of animals used are: Ctrl = 6, BKO = 5, Rh1-OE = 5, BKO-Rh1-OE =5. \**P*<0.05, \*\**P*<0.01 by Student’s *t*-test. Error bars represent s.e.m. (**C**) Representative images of ERα IHC in spontaneous mammary tumors. Each image represents an individual tumor. Scale bar= 50 μm. (**D**) Proposed model for the role of R-loops in *Brca1*-associated tumorigenesis.

Given that luminal progenitor cells are the cell of origin of *Brca1*-associated mammary tumors, we compared via flow cytometry the relative abundance of luminal progenitor cells in various mouse cohorts. Using CD49b as the established luminal progenitor marker (Shehata et al. 2012)(fig. S2A), we observed an increased luminal progenitor cell population in BKO versus Ctrl animals (Fig. 4B), which is consistent with the previous findings from us and others (Lim et al. 2009; Nair et al. 2016; Chiang et al. 2019). Intriguingly, RNase H1 overexpression in *Brca1*-deleted mammary glands (BKO-Rh1-OE) resulted in an appreciable shift from CD49b^+^ luminal progenitor to CD49b^-^ mature luminal cell populations, rendering the relative abundance of these two cell populations similar to what was observed in the Ctrl animals (Fig. 4B). Accordingly, while all mammary tumors from the BKO cohort were ERα^-^, a significant percentage of *Brca1*-associated mammary tumors with RNase H1 overexpression from the BKO-Rh1-OE cohort expressed ERα (Fig. 4C, fig. S5A and Table S1). These results suggest that R-loop removal may influence the *Brca1*-associated tumor subtype by reshaping the differentiation status of the cell of origin for *Brca1*-associated malignant transformation (Fig. 4D).

The aforementioned findings are consistent with our previously published *in vitro* work, which points to a role of R-loops in suppression of ERα-encoding *ESR1* transcription and impediment of luminal cell differentiation (Chiang et al. 2019). In particular, we found that *in vitro* RNase H1 overexpression in primary breast cells isolated from human *BRCA1* mutation carriers promotes the transition from luminal progenitor cells to mature luminal cells (Chiang et al. 2019). Thus, instead of directly promoting tumorigenesis, *Brca1*-associated R-loop accumulation likely contributes to luminal differentiation blockage and aberrant expansion of the luminal progenitor cell population in *Brca1*-deficient mammary glands. RNase H1 overexpression alleviates this differentiation roadblock and gives rise to more *Brca1*-deficient mature luminal cells, which, due to elevated genome instability, can undergo further oncogenic transformation into ERα^+^ tumors (Fig. 4D). Our findings provide *in vivo* evidence for a previously unappreciated role of R-loop dynamics on tumor development in mammary epithelium.

We envision that the conditional transgenic mouse model for RNase H1 overexpression provides a useful tool for probing the physiological impact of R-loop dynamics in other tissues and cell types. However, we also note several technical caveats and limitations of this transgenic model. First, a pattern of heterogeneous RNase H1 overexpression was observed in different mammary tumors from BKO-Rh1-OE mice (fig. S5B and table S1). In addition, given the myriad functions of R-loops in DNA transactions, global RNase H1 overexpression can reduce both beneficial and deleterious R-loops, thus complicating the interpretation of the corresponding *in vivo* phenotypes. Lastly, excessive RNase H1 may also give rise to “off-targeting” effects independent of R-loop removal. Despite these caveats, our current work fills an important gap in the knowledge of the pathophysiological roles of R-loops. By challenging the prevailing paradigm concerning the molecular functions of R-loops, our *in vivo* findings call for a more direct and rigorous examination of the presumed causal relationship between R-loops and tumorigenesis.

## Materials and Methods

### Mice

*Brca1^f/f^* and *MMTV-Cre* line A mice have been described previously (Zhang et al., 2017). *Rnaseh1^Tg^* conditional knockin mice were generated by CRISPR-based gene editing (Cyagen Biosciences). Briefly, the “CAG-*loxP*-Stop-*loxP-*mouse *Rnaseh1* cDNA-polyA” cassette was cloned into intron 1 of *Rosa26* in reverse direction. *Cas9* protein and gRNA were co-injected with donor vector into fertilized mouse eggs to generate the targeted knockin offspring. F0 founder animals, identified by PCR and subsequent sequence analysis, were bred to WT mice and assessed for germline transmission and F1 mouse generation. *MMTV-Cre* mice were used to generate *MMTV-Cre* (Ctrl)*, MMTV-Cre, Brca1^f/f^* (BKO), *MMTV-Cre, Rnaseh1^Tg/+^* (Rh1-OE) and *MMTV-Cre, Brca1^f/f^*, *Rnaseh1^Tg/+^* (BKO-Rh1-OE) mice. All mutant mice and their littermate controls were in a similarly mixed genetic background (129SvEv/SvJae/C57BL6/FVB). Genotyping primers information can be found in table S2. All procedures performed on animals were approved by the Institutional Animal Care and Use Committee at the George Washington University.

### Whole mount and immunohistochemistry staining

Inguinal mammary glands from mice of the indicated age were used for whole-mount staining as described previously (Nair et al., 2016). In brief, inguinal fat pads were isolated and spread onto a slide. Glands were fixed in Carnoy’s fixative overnight at room temperature. Glands were rehydrated in descending grades of ethanol (100, 70, 50 and 30%) for 15 min each, then washed with distilled water twice before overnight staining in Carmine alum solution (0.2% carmine and 0.5% aluminum potassium sulfate). Stained glands were dehydrated in ascending grades of alcohol (70, 70, 90, 95, 100 and 100%) for 15 min each, and put in CitriSolv reagent (Fisher, 22-143975) for tissue clearing. Samples were covered with sufficient Permount (Electron Microscopy Sciences, 1798605) for mounting. Samples were examined under a Nikon SMZ1000 dissection microscope. Duct length was measured from calibrated images using Eclipse software. Ductal fill percentage was calculated by the average length of three longest ducts originating from the nipple region divided by the overall fat pad length of each animal. Each dot represents a mammary fat pad from individual animals of a given genotype.

Mammary glands or mammary tumors were formalin-fixed overnight at 4 °C and paraffin embedded (FFPE). Sections of 3 μM in thickness were used for hematoxylin–eosin (H&E) staining, immunohistochemistry (IHC), and immunofluorescent (IF) staining. Sections were baked at 70 °C for 10 min, then de-paraffinized and dehydrated by xylene twice, and descending grade of ethanol (100 100, 95, 70 and 50%). Samples were washed briefly with PBS before transferring to boiling antigen-unmasking solution (Vector Labs, H-3300) for 20 min. For IHC, sections were pre-treated with 3% hydrogen peroxide for 10 min before blocking (10% normal goat serum in PBS) for 1 hr at room temperature. Samples were incubated with primary antibody in blocking buffer overnight at 4 °C. The ABC peroxidase detection system (Vector Labs, PK-6105) was used with 3, 3′-diaminobenzidine (DAB) (Vector Labs, SK-4105) as substrate according to the manufacturer’s instruction. Primary antibodies used were anti-RNase H1 (Proteintech, 15606-1-AP, 1:1000), anti-milk protein (Nordic Immunology, RAM/MSP, 1:10,000), and anti-ERα (Santa Cruz, sc-542, 1:500)

### R-loop IF staining

Freshly cut 3 μM FFPE samples were baked at 70 °C for 15 min. After de-paraffin and rehydration, samples were treated with boiling antigen-unmasking solution for 1 h. Samples were cooled down to room temperature, and then treated with 0.2 × SSC buffer (Ambion, AM9763) with gentle shaking at room temperature for 20 min. Samples were then incubated in a staining buffer (TBST with 0.1% BSA) for 10 min with rocking. Enzymatic treatments were done in staining buffer supplemented with 3 mM magnesium chloride with 1:200 dilutions of RNase T1 (Thermo Fisher Scientific, EN0541), ShortCut RNase III (New England Biolabs, M0245S), and/or RNase H (New England Biolabs, M0297S) and incubated with rocking for 1 hr. Samples were subsequently washed by incubating with staining buffer for 10 min with rocking. Primary antibody incubation was done with monoclonal antibody S9.6 (Karafast, ENH001) at 1:100 dilution in PBS containing 1% normal goat serum and 0.5% Tween-20 at 37 °C overnight. After primary antibody incubation, samples were washed three times with PBS containing 0.5% Tween-20, and then incubated with Alexa-488-conjugated secondary antibody at 1:1,000 dilution in PBS containing 1% normal goat serum and 0.5% Tween-20 at 37 °C for 2 hr. Samples were washed twice with PBS containing 0.5% Tween-20, twice with PBS, and then mounted with ProLong™ Gold Antifade Mountant with DAPI (Invitrogen, P36931). For each experiment, all samples were prepared, treated, and stained in parallel from one master enzyme, antibody, and/or dye dilution to ensure uniform treatment and staining efficiencies. R-loop intensity was quantified using the MetaMorph Microscopy Automation and Image Analysis Software 7.8 as previously described (Zhang et al., 2017).

### *In vivo* BrdU labeling and IF staining

For BrdU/RAD51 and BrdU/γΗ_2_ΑX double staining after IR, mice were first labeled with BrdU *in vivo*. Mice were intraperitoneally injected with cell proliferation labelling reagent (GE Healthcare, RPN201) at 16.7 ml kg^−1^ and then Gamma irradiated at 20 Gy. Three hours later, mammary glands were harvested for FFPE sectioning and IF staining. 3 μM sample sections were baked, de-paraffinized, rehydrated, and treated with boiling antigen-unmasking solution for 20 min. After cooling down, samples were washed with PBS 3 times and treated with 0.2% Triton X-100 for 30 min at room temperature. Samples were washed 3 times with PBS before putting in a blocking buffer for at least 1 hr. Samples were incubated with primary antibodies in blocking buffer overnight at 4 °C. Primary antibodies used were anti-BrdU (GE Healthcare, RPN20, 1:10,000), anti-γH_2_AX (Cell Signaling, 9718, 1:500), and anti-RAD51 (Santa Cruz, sc-8349, 1:100). The next day, sections were incubated with Alexa-488 and Alexa-546-conjugated secondary antibodies (Invitrogen, A32731 and A11126), mounted with ProLong™ Gold Antifade Mountant with DAPI, and examined with a Zeiss Spinning Disk Confocal Microscope.

### Flow cytometry and cell sorting

Thoracic and inguinal mammary glands from virgin mice were isolated in a sterile condition and lymph nodes from inguinal glands were removed. Single cells were prepared as previously described (Nair et al. 2016). Briefly, isolated glands were minced and digested for 16-20 hr at 37 °C in DMEM F-12 (StemCell Technologies, 36254) containing 2% fetal bovine serum (FBS), insulin (5 mg/mL), penicillin–streptomycin and 10% gentle collagenase/hyaluronidase (StemCell Technologies, 07919). After vortexing, epithelial organoids were pelleted at 600*g* for 5 min. Red blood cells (RBCs) were lysed with 0.8% ammonium chloride solution (StemCell Technologies, 07850). Epithelial organoids were further digested by 0.05% pre-warmed Trypsin (Life Technologies, 25300), washed in ice-cold Hanks Balanced Salt Solution (StemCell Technologies, 37150) with 2% FBS (HF), and resuspended in 5 U/mL dispase (StemCell Technologies, 07913) with 0.1 mg/mL DNase I (Sigma-Aldrich, D4513). Single cells were obtained by filtering the cell suspension through a 40-μm cell strainer (Fisher, 22363547). Cells were counted and resuspended in HF at a concentration of 1 × 10^6^ cells per 100 μl for staining. Cell were first blocked with 10% rat serum (Jackson Laboratories, 012-000-120) for 10 min followed by incubation with cell-surface antibody cocktails for 20 min at 4 °C. The following cell-surface markers were used in the experiment: EpCAM-PE (BioLegend, 118206, 1:200), CD49f-FITC (BD Biosciences, 555735, 1:50), CD31-Biotin (BD Bioscience, 553371, 1:100), CD45-Biotin (BioLegend, 103103, 1:100), TER119-Biotin (BioLegend, 103511, 1:100), and CD49b-Alexa Fluor 647 (BioLegend, 104317, 1:200) followed by Streptavidin-Pacific Blue (Invitrogen, S11222, 1:200) incubation. 7-AAD (BD Biosciences, 559925, 1:20) was added 10 min before analysis. CD49b^+^ cells were gated using a fluorescent-minus-one control, in which all antibodies except CD49b-Alexa 647 were used. Flow cytometry was performed with a BD Celesta Cell Analyzer and sorting was performed with a BD Influx Cell Sorter. Data were analyzed using FlowJo software. Purity of the stromal, luminal, and basal populations were verified by RT-PCR analysis of *Vimentin* (stromal), *K18* (luminal), and *K14* (basal) mRNA. All primers information can be found in table S2.

### Statistical analysis

Data analysis and statistics were done using GraphPad Prism 8 software. Mean difference comparison from two groups using unpaired student *t*-test was used throughout the experiments. Data in bar and dot graphs are means ± s.e.m. *P* < 0.05 was considered statistically significant.

### Data and materials availability

All data needed to evaluate the conclusions in the paper are present in the paper and the Supplementary Materials. Model demonstration is created with BioRender.com.

## Acknowledgments

We thank Madeleine Prevost, Shannen Ubalde, and Yimeng Huang for technical support.

## Author contributions

R.L. and Y.H. conceived, managed, and oversaw the overall project. R.L. and H-C.C. wrote the manuscript. H-C.C., L.Q., and P.M. carried out the experiments. H-C.C., L.Q., P.M., Y.H., and R.L. analyzed the data.

